# Rational design of structure-based vaccines targeting misfolded alpha-synuclein conformers of Parkinson’s disease and related disorders

**DOI:** 10.1101/2023.06.30.547254

**Authors:** Jose Miguel Flores-Fernandez, Verena Pesch, Aishwarya Sriraman, Enrique Chimal-Juarez, Sara Amidian, Xiongyao Wang, Sara Reithofer, Liang Ma, Gültekin Tamgüney, Holger Wille

**Author notes:** These authors contributed equally to this work. Deceased. Current addresses: Indiana University School of Medicine, Stark Neurosciences Research Institute, Indianapolis, IN, USA. School of Materials Science and Engineering, Harbin Institute of Technology, Weihai, Shandong, China (Xiongyao Wang). Corresponding authors: - Gültekin Tamgüney & - Holger Wille.

## Abstract

**Background:** Synucleinopathies, including Parkinson’s disease, multiple system atrophy, and dementia with Lewy bodies, are neurodegenerative disorders caused by the accumulation of misfolded alpha-synuclein protein. Developing effective vaccines against synucleinopathies has been challenging due to the difficulty of stimulating an immune-specific response against alpha-synuclein, conferring neuroprotection without causing harmful autoimmune reactions, and selectively targeting only pathological forms of alpha-synuclein. Previous attempts using linear peptides and epitopes without control of the antigen structure for immunization failed in clinical trials. The immune system was unable to distinguish between the native alpha-synuclein and its amyloid form.

**Results:** The prion domain of the fungal HET-s protein was selected as a scaffold to introduce select epitopes from the surface of alpha-synuclein fibrils. Four vaccine candidates were generated by introducing specific amino acid substitutions onto the surface of the scaffold protein in regions that showed structural similarity to alpha-synuclein fibril structures. Each vaccine candidate had unique amino acid substitutions that imitated a specific epitope from alpha-synuclein amyloid fibrils. The approach successfully mimicked the stacking of the parallel in-register beta-sheet structure seen in alpha-synuclein fibrils as the vaccine candidates were found to be structurally stable and self-assembling into the desired conformations. All vaccine candidates induced substantial levels of IgG antibodies that recognized pathological alpha-synuclein fibrils derived from a synucleinopathy mouse model. Furthermore, the resulting anti-sera recognized pathological alpha-synuclein aggregates in brain lysates from patients who died from dementia with Lewy bodies, multiple system atrophy, or Parkinson’s disease, but did not recognize linear alpha-synuclein peptides. Each vaccine candidate induced a unique pattern of reactivity toward alpha-synuclein aggregates contained in distinct disease pathologies.

**Conclusions:** This new approach, based on the rational design of vaccines using the secondary and tertiary structure of alpha-synuclein amyloid fibrils and strict control over the exposed antigen structure used for immunization, as well as the ability to mimic aggregated alpha-synuclein, provides a promising avenue towards developing effective vaccines against alpha-synuclein fibrils, which may be crucial for the prevention and treatment of synucleinopathies.

## Background

Synucleinopathies are a group of neurodegenerative disorders characterized by the abnormal accumulation of misfolded alpha-synuclein in the brain [1]. The disorders include among others Parkinson’s disease (PD), dementia with Lewy bodies (DLB), multiple system atrophy (MSA), progressive supranuclear palsy (PSP), and the Lewy body variant of Alzheimer’s disease [1–4].

Prevalence and mortality rates for the synucleinopathies, including Parkinson’s disease, vary across regions and studies. Parkinson’s disease is the second most common progressive neurodegenerative disorder, affecting 1-3% of people aged 65 or older [5]. Its prevalence, disability burden, and mortality rate are increasing faster than any other neurological disorder worldwide [6,7]. The number of Parkinson’s disease patients has more than doubled from 2.5 to 6.1 million between 1990 and 2016 globally, and this increase is not solely attributable to an aging population but also to other contributing factors, such as greater exposure to risk factors and longer disease duration [8]. Additionally, the number of people with dementia worldwide is expected to reach 153 million by 2050, indicating a 166% increase from the cases reported in 2019 [9].

Synucleinopathies share similar motor and non-motor symptoms such as tremors, muscle rigidity, bradykinesia, dyskinesia, instability, difficulties with balance and coordination, and often include cognitive impairment [4,10–14]. However, each disease has differing motor and non-motor symptoms that help distinguish them from one another. For example, Parkinson’s disease is characterized by an asymmetrical onset of symptoms and resting tremors [4], while dementia with Lewy bodies presents with fluctuating cognition and visual hallucinations [11]. Conversely, multiple system atrophy is marked by cerebellar ataxia and autonomic dysfunction [10]. It is thought that those symptoms are linked by the region of the brain where the abnormal accumulation of alpha-synuclein takes place. In dementia with Lewy bodies, the anomalous accumulation of alpha-synuclein appears to be more prevalent in limbic and neocortical regions, particularly in the temporal lobe and the CA2 area of the hippocampus, as compared to Parkinson’s disease [15]. Conversely, Parkinson’s disease cases show higher levels of dopaminergic cell loss in the substantia nigra, mainly affecting the dorsolateral regions, compared to dementia with Lewy bodies [15,16], while in multiple system atrophy, the accumulation of alpha-synuclein is found in oligodendrocytes in white matter tracts [17,18]. Although there may be some overlap in the distribution patterns of alpha-synuclein aggregates and symptoms among these diseases, each also presents unique motor and non-motor symptoms that are used to distinguish them.

Alpha-synuclein is an acidic protein of 140 amino acids that N-terminally folds into an α-helical structure when bound to a membrane [19]. In vivo, this protein is primarily found in a soluble state, particularly in the presynaptic terminals of neurons, and its precise function is not fully understood, but it is thought to play a role in regulating neurotransmitter release and synaptic plasticity to allow communication between nerve cells [20]. Nonetheless, alpha-synuclein can convert into insoluble, structured amyloid fibrils. These fibrils contain a characteristic beta-sheet structure [21,22], forming fibrillar deposits termed Lewy bodies and Lewy neurites, which are hallmarks in the pathogenesis of synucleinopathies. Although the precise mechanisms by which alpha-synuclein converts into amyloid fibrils are not yet fully understood, the core region of alpha-synuclein, which consists of residues 61-95, has been identified as a key determinant in the aggregation and fibril formation of the protein [23]. The misfolding and subsequent aggregation of alpha-synuclein can propagate via intercellular and synaptic transmission, invading multiple regions of the brain and producing cytotoxic effects that cause the progressive loss of dopaminergic neurons (50-70%) in the substantia nigra pars compacta, leading to the onset of motor symptoms in Parkinson’s disease [24,25].

Prevention and effective therapies are ongoing research priorities for all synucleinopathies and Parkinson’s disease in particular. However, to date, no neuroprotective, restorative, or disease-modifying therapy is available [26]. Currently available treatment options are symptomatic only, designed to replace dopamine function without any neuroprotective activity [27]. In addition, it has been shown that prolonged use of therapeutic drugs, such as levodopa and amantadine, often has variable therapeutic effects and leads to the development of undesirable adverse reactions, besides affecting cytokine production, suggesting that not only PD itself but also the therapeutic approach may affect patients’ immune systems [28]. Ongoing research in the field of synucleinopathies aims to prevent or halt disease progression, with various approaches under investigation, including immunotherapy, gene therapy, stem cell therapy, and targeting alpha-synuclein aggregation.

In the case of Parkinson’s disease, one promising avenue is to prevent the formation of amyloid fibrils by targeting alpha-synuclein. This can be achieved through passive and active immunization approaches, as well as small molecule inhibitors with neuroprotective effects. Passive immunization involves injecting preformed (monoclonal) antibodies that target and clear or reduce the accumulation of alpha-synuclein aggregates from the brain, while active immunization involves administering vaccines that trigger an immune-specific response against alpha-synuclein to prevent or reduce the accumulation of aggregates [28]. However, if an effective passive immunotherapy to stop PD progression were found, repeated injections would be required, resulting in high costs, and this approach may not be beneficial in the long term, as alpha-synuclein amyloid fibrils may continue to accumulate, and chronic neuroinflammation would persist [28,31,32].

Here, nanobiotechnology can enable the improvement of antigen delivery to the immune system and the enhancement of immune responses, making it a valuable tool for developing vaccines and immunosuppressants, as it has been shown to stimulate local cytokine production, induce mucosal antibody responses, elicit different T cell responses, and aid in vaccine design targeting B-cells [33,34]. By producing and using materials in the nanoscale range, researchers can take advantage of unique size- and structure-dependent properties and phenomena that occur at this scale, allowing for the development of targeted immunotherapies for diseases such as Parkinson’s disease.

One of the challenges in developing active immunization therapies has been to create a vaccine that can stimulate an immune response against alpha-synuclein, confer neuroprotection without causing harmful autoimmune reactions, and selectively target only the pathological forms of alpha-synuclein fibrils while leaving the physiologically active form intact. Although these challenges have not yet been overcome, clinical trials are currently underway to test the safety and efficacy of vaccines targeting synucleinopathies in humans [35–37]. However, attempts to develop vaccines for Parkinson’s disease and other synucleinopathies have failed because most studies have used the native form of alpha-synuclein rather than its amyloid form, which the immune system is unable to distinguish from the pathological form [36,38–44].

To overcome this challenge, it is crucial to identify specific antigenic targets on the surface of the pathological forms of alpha-synuclein, which can be targeted by the immune system. These targets could include specific regions of the protein that are only exposed in the pathological forms, or post-translational modifications that are unique to the pathological forms. Therefore, the goal of this study is to develop effective active immunotherapy agents for Parkinson’s disease and other synucleinopathies by designing vaccines that mimic regions of the alpha-synuclein protein that are surface exposed in its aggregated/amyloid form only. In contrast to the conventional use of peptide antigens and linear epitopes, this study presents a novel approach based on antigens that mimic small, specific surface structures found on amyloid fibrils of aggregated alpha-synuclein protein. These antigens are based on structural carriers that are engineered to include carefully selected surface residues, allowing control over the conformation of the resulting epitope and generating a specific immune response, resulting in an effective active immunization as prophylaxis to prevent disease. By using this approach, we aim to induce an immune response against pathologic alpha-synuclein species that can have a significant protective and, potentially, also a therapeutic benefit for patients with Parkinson’s disease, dementia with Lewy bodies, multiple system atrophy, and other synucleinopathies.

## Results

### Structural basis for the design of alpha-synuclein-targeting vaccine candidates

To mimic the structural features of alpha-synuclein amyloid fibrils and design vaccine candidates targeting synucleinopathies, a scaffold protein was selected to introduce exposed amino acids from the surface of alpha-synuclein fibril structures in a structurally controlled manner. The innocuous HET-s prion domain from the fungus *Podospora anserina*, which has a left-handed, two-rung beta-solenoid fold structure with a distance of 4.8 ± 0.2 Å between each beta-rung, and containing three beta-strands per rung [45,46], was chosen as the scaffold protein for this purpose. The HET-s prion domain HET-s(218-289) was selected due to its innocuous nature, well-defined structure, propensity to fold into a unique conformation *in vitro*, and a rational analysis showing structural similarities to alpha-synuclein fibrils based on similar structural arrangements of the cross-beta elements (Fig. 1).

**Figure 1:**
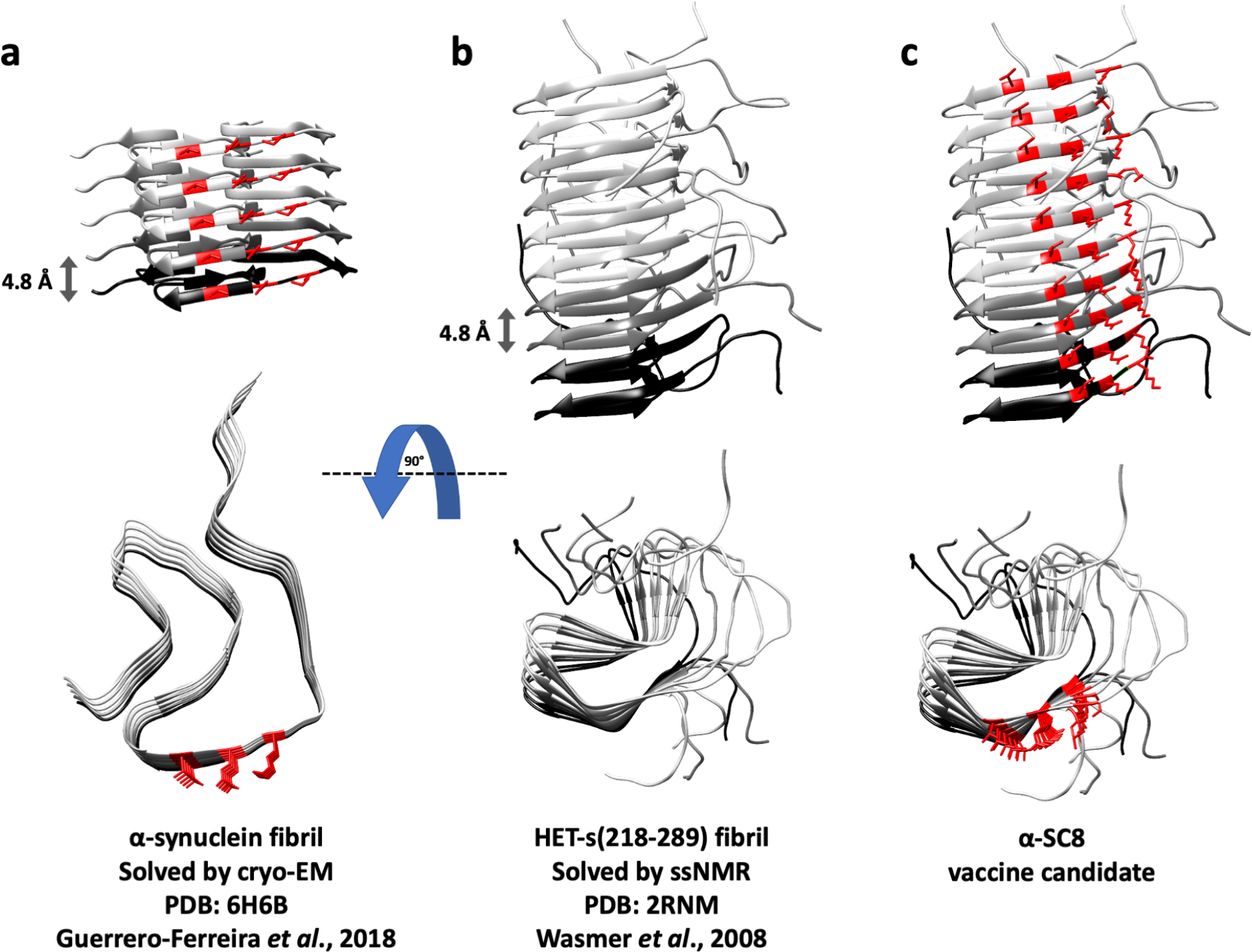
Rational design approach for developing structure-based vaccines that target misfolded alpha-synuclein conformers associated to Parkinson’s disease and related disorders at the structural level. The approach is based on similarities of cross-beta structural arrangements and the cross-sectional views of a) an alpha-synuclein fibril showing a parallel in-register beta-sheet structure with 8 to 12 beta-strands and a characteristic spacing of ∼4.8 Å between beta-strands, and b) the HET-s(218-289) fibril structure, showing a left-handed, two-rung beta-solenoid fold with three beta-strands per rung and a distance of ∼4.8 Å between each beta-rung. c) Schematic representation of a rationally designed vaccine candidate for a synucleinopathy disorder, which involves introducing exposed amino acids from the surface of the alpha-synuclein fibril structure using HET-s(218-289) as a scaffold protein.

The overall structural similarity between HET-s(218-289) and alpha-synuclein allowed for the introduction of exposed amino acids from the surface of alpha-synuclein fibril structures in a structurally controlled manner. This was possible because the three-dimensional structures of alpha-synuclein amyloid fibrils had been deciphered, revealing a fibril core consisting of 8 to 12 beta-strands forming a parallel in-register beta-sheet structure with a characteristic spacing of ∼4.8 Å between beta-strands. The number and position of the beta-strands varies depending on the strain or conformer of alpha-synuclein aggregates, which can be generated *in vitro* (Figs. 1 and 2 a,c) or isolated *ex vivo* from the brains of individuals affected by multiple system atrophy, dementia with Lewy bodies, or Parkinson’s disease [47–50].

**Figure 2:**
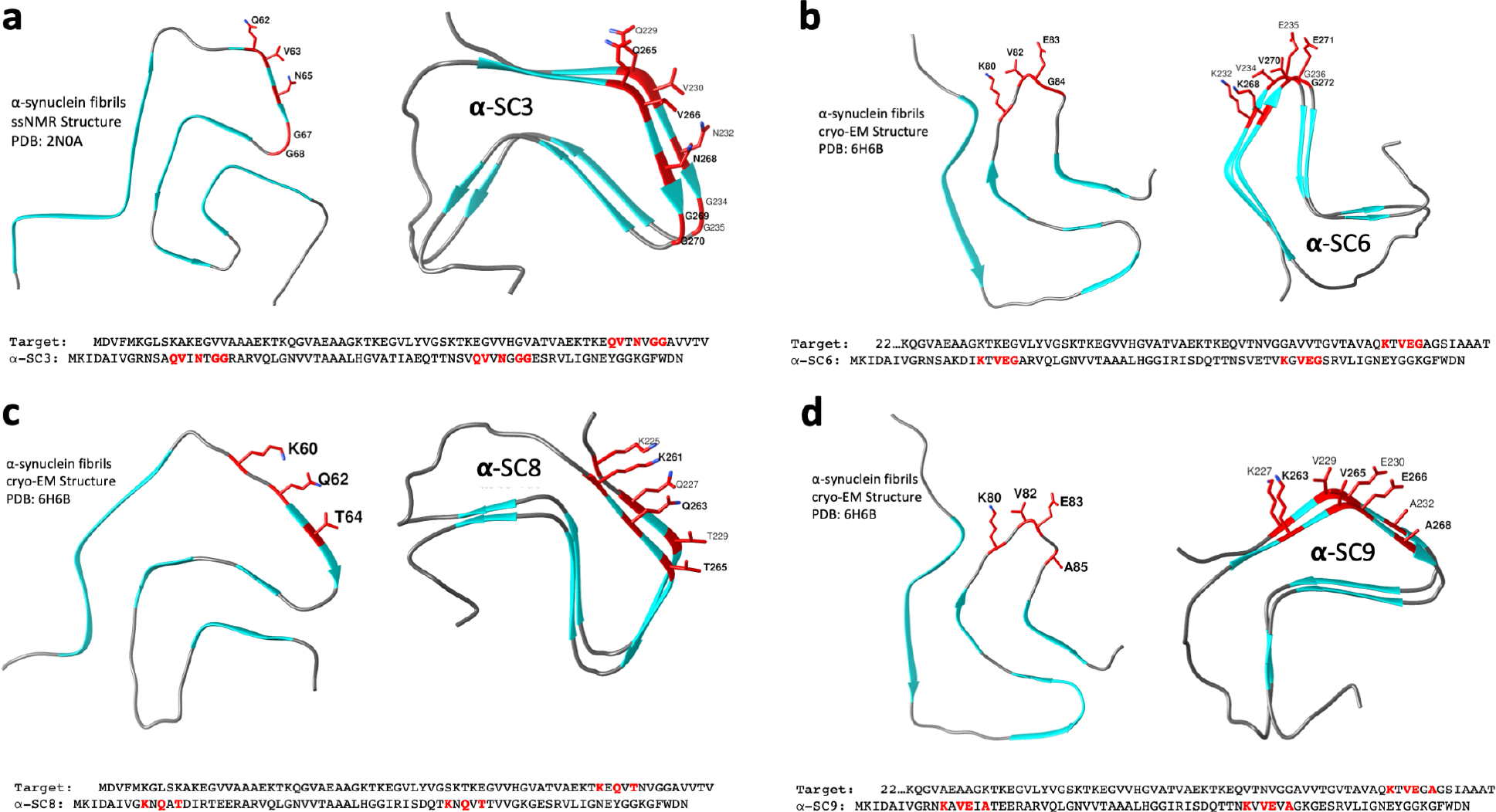
Structural basis of the four engineered vaccine candidates, α-SC3, α-SC6, α-SC8, and α-SC9, designed to target alpha-synuclein fibrils. The HET-s(218-289) prion domain was used as a scaffold to mimic select epitopes from the surface of two different alpha-synuclein amyloid fibril structures to encompass fibrillar heterogeneity. Specific amino acid substitutions (highlighted in red) were introduced onto the surface of both rungs of the beta-solenoid scaffold protein in regions that showed structural similarity to the alpha-synuclein fibril structures to mimic the stacking of two alpha-synuclein molecules per two rungs of the scaffold protein. For the vaccine candidate a) α-SC3, the amino acids substitutions (K229Q, D230V, R232N, E234G, and E235G) were introduced on the first rung of the scaffold protein, and (E265Q, T266V, V268N, and K269G) on the second rung, while for b) α-SC6, the substitutions (R232K, E234V, and R236G) were introduced on the first rung and (V268K, K270V, G271E, and E272G) on the second rung. For c) α-SC8, the substitutions (R225K, S227Q, and K229T) were introduced on the first rung and (T261K, S263Q, and E265T) on the second rung. Finally, for d) α-SC9, the substitutions (S227K, K229V, D230E, and R232A) were introduced on the first rung and (S263K, E265V, T266E, and V268A) on the second rung.

### Design and engineering of alpha-synuclein-targeting vaccine candidates

To design vaccines targeting aggregated alpha-synuclein, we engineered the prion domain of the HET-s protein, HET-s(218-289) as an innocuous scaffold protein to mimic select epitopes from the surface of alpha-synuclein amyloid fibrils. Specifically, we introduced select amino acids from the surface of alpha-synuclein fibril structures (Fig. 2) into the scaffold protein in a structurally controlled, discontinuous manner to express specific antigenic determinants. We used two different alpha-synuclein structures to encompass fibrillar heterogeneity, one solved by Tuttle et al. [50] using solid-state NMR and the other by Guerrero-Ferreira et al. [49] using cryo electron microscopy.

Based on the analyses, we successfully engineered four different vaccine candidates named α-SC3 (Fig. 2a), α-SC6 (Fig. 2b), α-SC8 (Fig. 2c), and α-SC9 (Fig. 2d) from an initial collection of ten designs (data not shown). Each candidate was generated by introducing specific amino acid substitutions onto the surface of the scaffold protein in regions that showed structural similarity to the alpha-synuclein fibril structures. The substitutions of residues were carefully selected to mimic the structure of two alpha-synuclein molecules per rung of the beta-solenoid scaffold. Thus, the repeating nature of the HET-s(218-289) beta-solenoid assembly effectively mimicked the stacking of the parallel in-register beta-sheet structure of the alpha-synuclein fibrils (e.g. Fig. 1).

For instance, α-SC3 included substitutions of K229Q, D230V, R232N, E234G, and E235G for the first rung of the scaffold protein to mimic a molecule of alpha-synuclein in its amyloid form, while the residues E265Q, T266V, V268N, and K269G were replaced in the second rung of the scaffold protein to mimic a second molecule of alpha-synuclein that would sit on top of the first one in its amyloid form (Fig. 2a). Similarly, for the other vaccine candidates, we carefully selected and substituted specific residues for each rung of the scaffold protein to mimic the structure of alpha-synuclein molecules in their amyloid form. Specifically, for α-SC6, we substituted R232K, E234V, and R236G for the first rung and V268K, K270V, G271E, and E272G for the second rung (Fig. 2b); for α-SC8, we substituted R225K, S227Q, and K229T for the first rung and T261K, S263Q, and E265T for the second rung (Fig. 2c); and for α-SC9, we substituted S227K, K229V, D230E, and R232A for the first rung and S263K, E265V, T266E, and V268A for the second rung (Fig. 2d).

It is worth highlighting that the HET-s prion domain has a two-rung beta-solenoid structure in its native amyloid fold. During the formation of its amyloid state, the next molecule stacks on top of the previous one. However, analyzing the structure in detail, it resembles the parallel in-register beta-sheet structure of alpha-synuclein amyloid fibrils by its surface exposure of alternating residues in the beta-strands (Fig. 1). By introducing specific amino acid substitutions into both rungs of the HET-s(218-289) scaffold, we were able to mimic the surface features of the pathological alpha-synuclein amyloid fibrils. Notably, when the modified HET-s fibrils form containing these specific substitutions, all rungs of the scaffold protein carry the same substitutions, resulting in a highly ordered and structurally homogeneous fibril. To test our hypothesis and the vaccine candidates, the coding sequences of the engineered proteins were optimized for expression in *Escherichia coli*, cloned into a pET-21a(+) expression vector, and inserted into *E. coli* BL21 (DE3) chemically competent cells. Once expressed, purified by affinity chromatography from inclusion bodies, and desalted (Fig. S1), the alpha-synuclein-targeting vaccine candidates underwent an *in-vitro* fibrillization process by neutralizing the pH and stirring the protein solutions at room temperature.

### Evaluation of the structural stability and self-assembly of the vaccine candidates

The structure-based rational design of the vaccines targeting misfolded alpha-synuclein did not affect the stability of the four proteins α-SC3, α-SC6, α-SC8, and α-SC9, as their secondary structures including the substituted residues were stable (Fig. 3). This was demonstrated by far-UV circular dichroism spectra, which showed a single minimum at around 220 nm, typical for beta-sheet proteins with negligible α-helical content (Fig. 3) [51]. However, in other constructs from the initial collection of ten designs, the substituted residues affected the structural integrity of the protein scaffold, as evidenced by CD profiles showing typical indications of random coil structure (data not shown).

**Figure 3:**
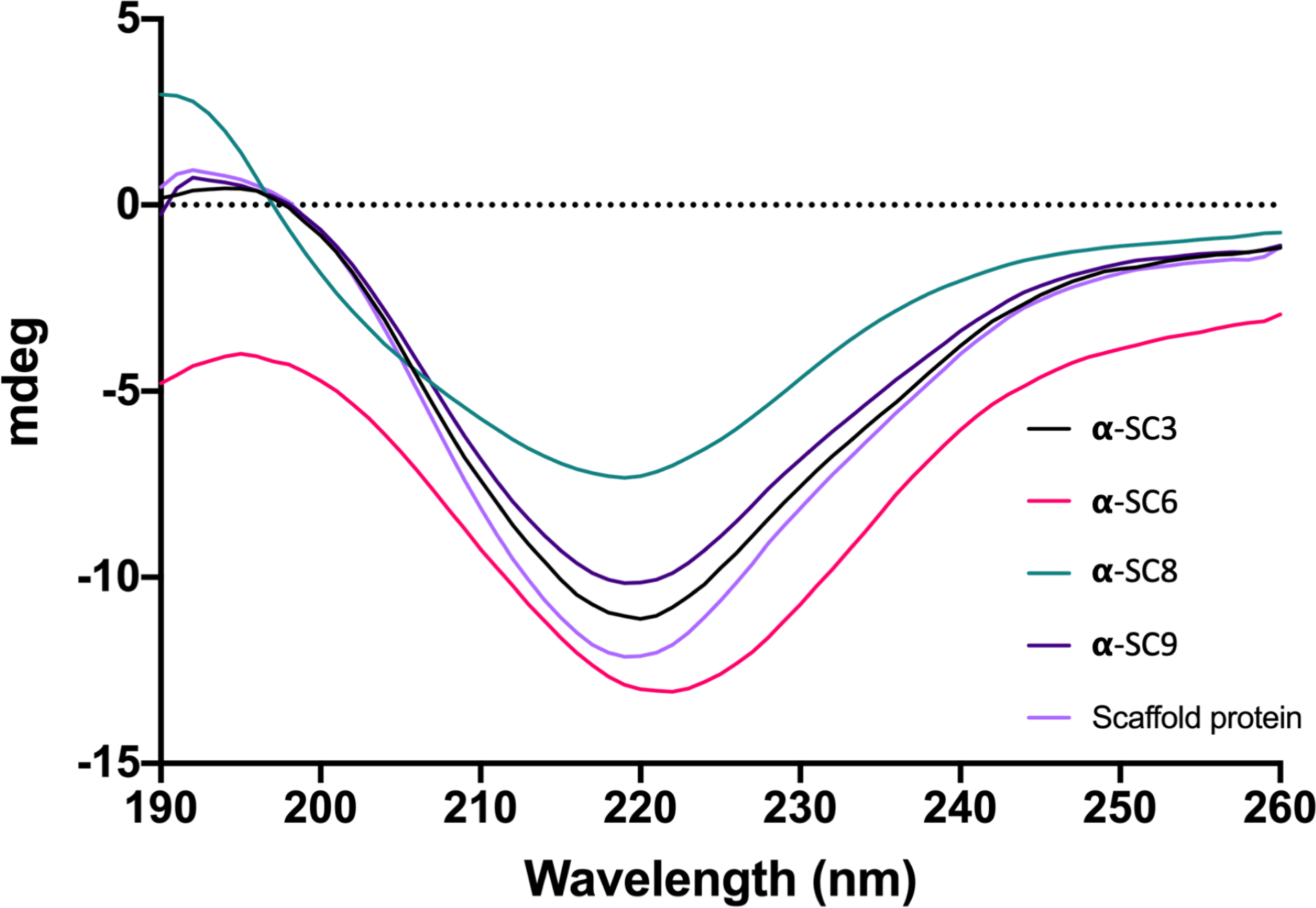
The far-UV circular dichroism spectra revealed a predominant beta-sheet secondary structure of the four engineered vaccine candidates (α-SC3, α-SC6, α-SC8, and α-SC9) designed to target misfolded alpha-synuclein, with a single minimum at ∼220 nm, indicative of a predominantly beta-sheet secondary-structure content. The CD profiles showed negligible alpha-helical content. The stability of the vaccine candidates and the scaffold protein HET-s(218-289) was evaluated, and the structural integrity of both was maintained after the introduction of specific amino acid substitutions onto the surface of both rungs of the beta-solenoid scaffold protein.

To confirm the ability of the vaccine candidates to self-assemble into the typical amyloid structure of the native HET-s(218-289) protein, negative stain transmission electron microscopy was used. The negative stain electron micrographs demonstrated the capability of the afore mentioned vaccine candidates to fibrillize into amyloid fibrils (Fig. 4a-d) that were indistinguishable by negative-stain transmission electron microscopy from those of the scaffold protein (Fig. 4e). This demonstrated that the constructs adopted the intended fold and were capable of self-assembly into the desired amyloid fibril structures. The next step in the development of these vaccine candidates was to evaluate their immunogenicity in a rodent model.

**Figure 4:**
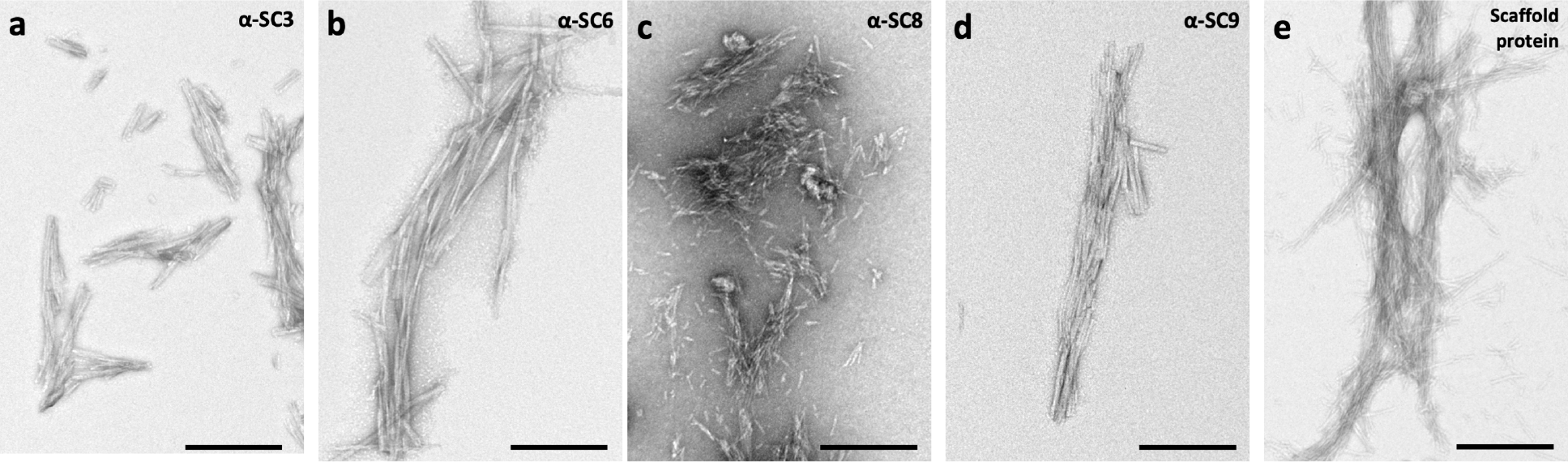
Negative stain transmission electron microscopy analysis demonstrated that the vaccine candidates adopted the intended fold and were capable of self-assembly into the desired amyloid fibril structures. The micrographs for a) α-SC3, b) α-SC6, c) α-SC8, and d) α-SC9 showed that the four vaccine candidates self-assemble into typical amyloid fibrils that are indistinguishable from those of the e) scaffold protein. Scale bar, 200 nm.

### Optimization of the immunization regimen for inducing high levels of IgG antibodies

To assess the immunogenicity of the vaccine candidates that adopted the proper beta-solenoid fold, mice were immunized intraperitoneally using a regime involving a priming dose of the vaccine candidate with Freund’s complete adjuvant, followed by three biweekly boosters of half the priming dose along with Freund’s incomplete adjuvant. To determine an optimal immunization dose, initial priming doses of 100, 50, 25, 5 µg of vaccine candidate α-SC9 were tested, and the titer of post immune sera was determined (Fig. 5a). The analysis revealed that the optimal dose of vaccine candidate was 25 µg for the initial priming dose and 12.5 µg for each booster dose.

**Figure 5:**
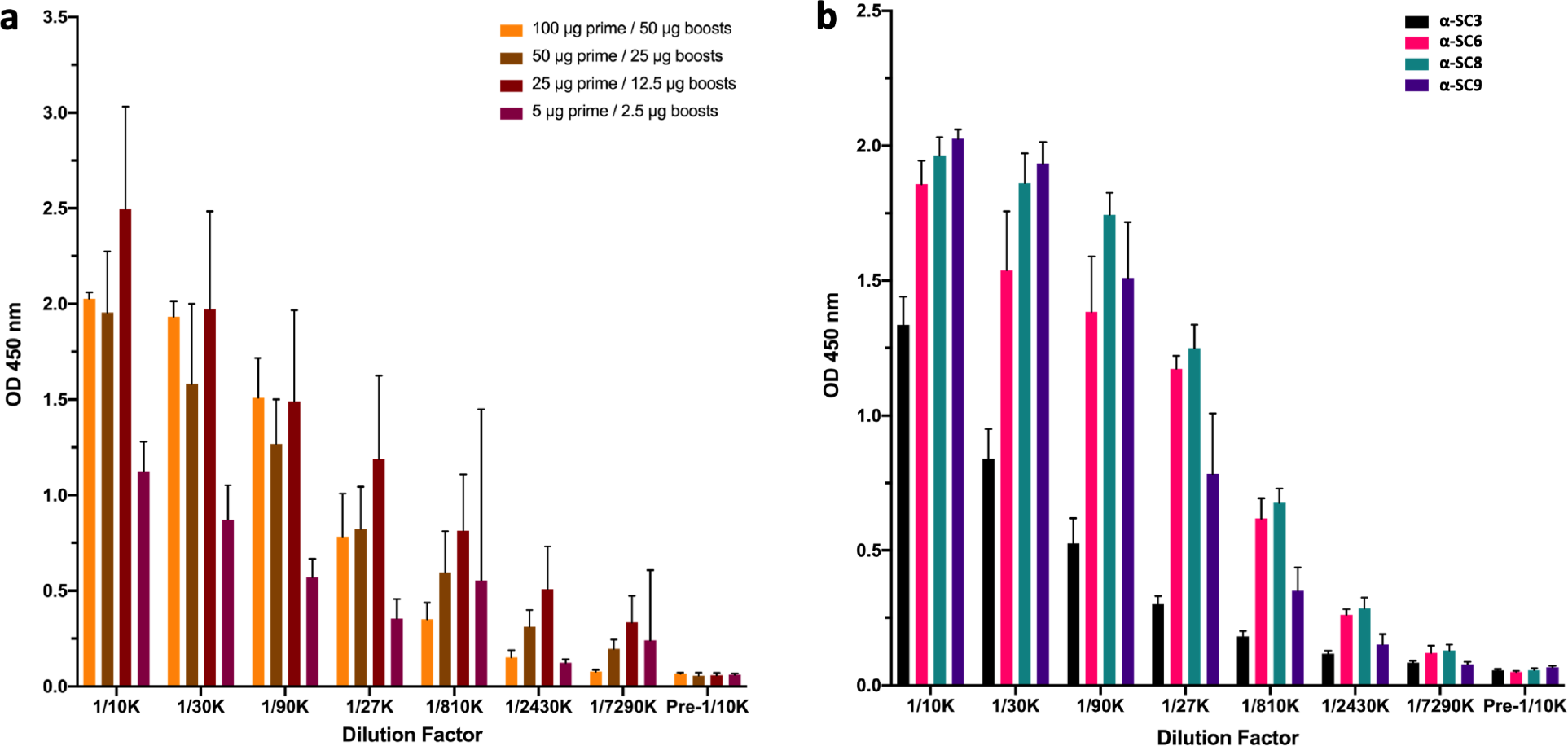
The vaccine candidates elicited specific immunoglobulin G antibody responses in the immunized FVB mice, as determined by ELISA. The optimal immunization dose for a) vaccine candidate α-SC9 was determined by administering initial priming doses of 100, 50, 25, and 5 µg, followed by three biweekly boosters of half the priming dose. Post-immune serum titers were determined for each dose. b) All mice were immunized with an initial dose of 25 µg followed by a booster dose of 12.5 µg. Pre-immune and post-immune antisera were analyzed by ELISA to detect specific IgG antibody responses. The data represent the mean value of four mice ± SEM. All vaccine candidates induced substantial levels of IgG antibodies compared to pre-immune antisera.

The recommended protein amount for immunizing mice is typically based on soluble proteins. However, in the present study, the vaccine candidates are in an insoluble amyloid form, making it crucial to determine the optimal dosage. Antigen solubility can affect its processing and presentation to the immune system, which can impact the strength and specificity of the resulting immune response. Soluble proteins are usually more efficiently processed by antigen-presenting cells, such as dendritic cells, resulting in a robust and specific immune response, while insoluble proteins may require additional processing steps or specialized antigen-presenting cells to elicit an immune response.

However, as is the case in the present study where the insoluble protein is a structural component of a pathogen or cell, using the insoluble form of the protein may more closely mimic the natural presentation of the antigen and elicit a more physiologically relevant immune response. Additionally, using insoluble proteins may activate different immune pathways or induce different types of immune cells, which can be desirable in certain settings [52,53].

These concentrations and dosage regime were followed for the rest of the vaccine candidates (Fig. 5b). It was observed that all vaccine candidates induced substantial levels of IgG antibodies when compared to the pre-immune sera. The next step was to investigate whether those vaccine candidates were capable of eliciting an immune response that can specifically target pathological alpha-synuclein in its amyloid/aggregated form.

### Vaccine candidate-derived antisera recognize alpha-synuclein in a synucleinopathy mouse model

The potential of the candidate vaccines to stimulate an immune response in mice could pave the way for the development of an effective immunotherapy for Parkinson’s disease and other synucleinopathies as they were engineered to mimic the amyloid fibril structure of alpha-synuclein, the misfolded protein implicated in the pathogenesis of these neurodegenerative disorders. An effective active immunotherapy aims to stimulate the body’s immune system to target specific antigens, such as misfolded alpha-synuclein, which could potentially slow or halt disease progression.

Hemizygous TgM83^+/−^ (B6;C3-Tg(*Prnp*-SNCA*A53T)83Vle/J) mice overexpress mutant human A53T alpha-synuclein and are widely used in Parkinson’s disease and synucleinopathy research due to the significantly elevated amounts of aggregated alpha-synuclein in their brain upon infection with pre-formed alpha-synuclein fibrils. The injection leads to the development of synucleinopathy symptoms after an incubation period of 3-4 months when injected intracranially, and 7.5 and 9.5 months when injected via intraperitoneal or intraglossal inoculation, respectively [54–56]. Therefore, a successful immune response by the vaccine candidates that recognizes or binds to aggregated/amyloid alpha-synuclein in the hemizygous TgM83^+/−^-mouse brain could have important implications for the development of new immunotherapies for synucleinopathies.

The ability of the post immune antisera to recognize pathological alpha-synuclein fibrils was assessed by testing the antisera with brain homogenates from TgM83^+/−^ mice were that intracerebrally injected with pre-formed alpha-synuclein fibrils to induce a synucleinopathy, and with a control group of TgM83^+/−^ mice that were intracerebrally injected with bovine serum albumin (BSA) only. The data indicate that besides recognizing the original antigens (Fig. 5b), the vaccine candidates can also recognize pathological alpha-synuclein fibrils, as evidenced by a significant difference [<0.001] in antisera response compared to brain lysate injected with BSA (Fig. 6), which does not cause neurological disease or neuropathology [56]. These vaccine candidates have demonstrated the potential to elicit an immune-specific response against alpha-synuclein fibrils derived from a mouse model of synucleinopathy, supporting our novel approach for vaccine development targeting Parkinson’s disease and other synucleinophaties (Fig. 6). In contrast, past strategies to develop vaccines for synucleinopathies have utilized linear peptides or linear epitopes that merely mimic alpha-synulcein in its native form [36,38–44], lacking control over the structure of the antigen used for immunization and failing to imitate pathological alpha-synuclein in its aggregated/amyloid form.

**Figure 6:**
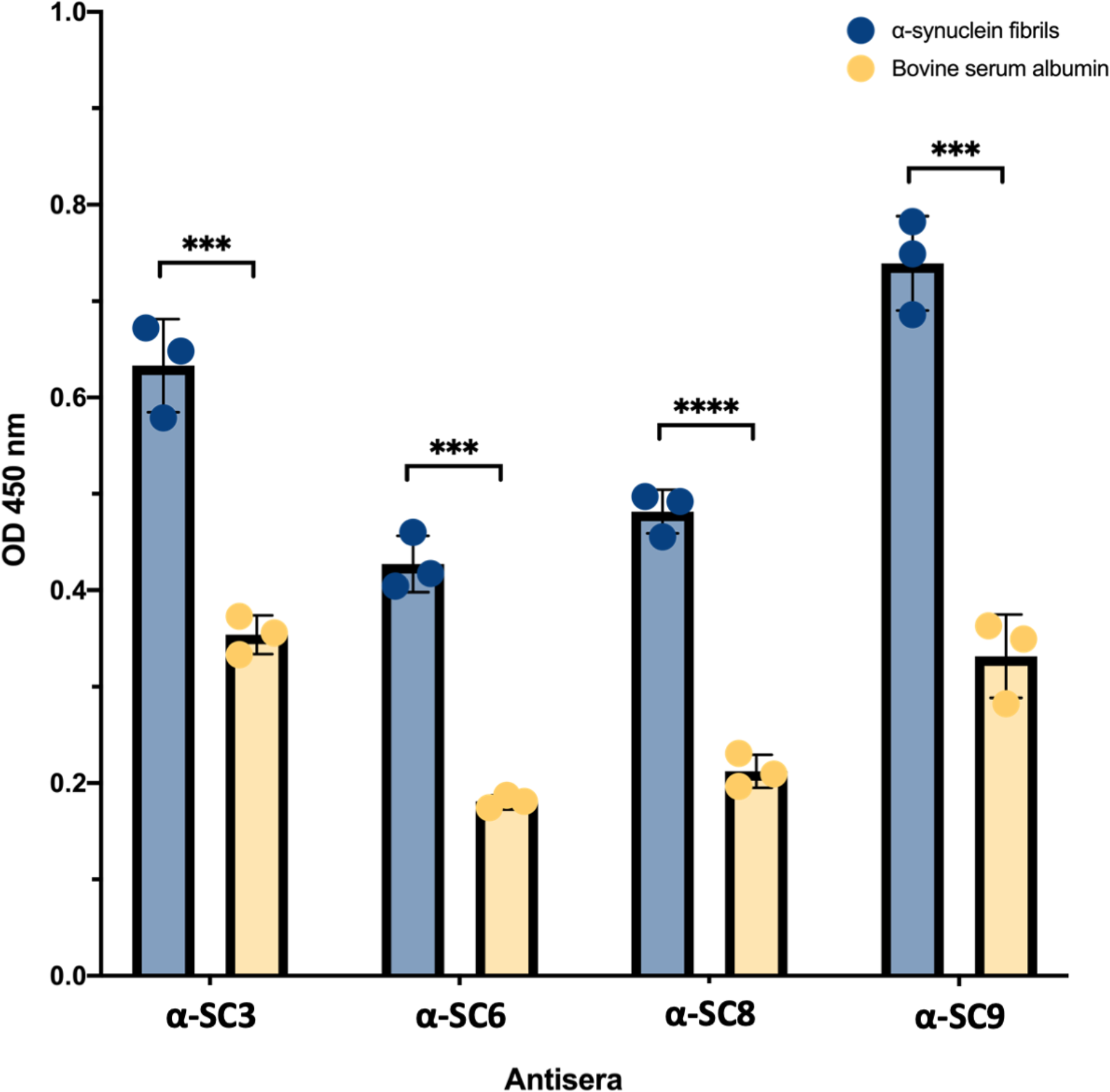
Detection of pathological alpha-synuclein fibrils from a synucleinopathy mouse model. Hemizygous TgM83^+/−^ mice were intracerebrally injected with preformed alpha-synuclein fibrils to induce a synucleinopathy, or with BSA as a negative control. Post-immune antisera of vaccinated wild-type mice recognized pathological alpha-synuclein fibrils in brain homogenate of diseased TgM83^+/−^ mice. This shows that the vaccine candidates elicited a significant immune-specific response against alpha-synuclein fibrils (p<0.001), demonstrating the potential of amyloid fibril-based vaccine development for Parkinson’s disease and other synucleinopathies. The data represent the mean value of four mice ± SEM. Statistical analysis was performed using a two-tailed paired t-Test [*** < 0.001, **** < 0.0001].

### Vaccine candidates induce antisera that recognize conformational but not linear alpha-synuclein epitopes

To further characterize and prove the ability of the post immune antisera elicited from the vaccine candidates to recognize the structure of alpha-synuclein fibrils instead of linear peptides/epitopes of alpha-synuclein in their native form, we tested the antisera with human brain lysates from patients who died with dementia with Lewy bodies, multiple system atrophy or Parkinson’s disease, and non-neurologic controls (Table 1), as well as with a set of linear peptides that comprised the entire sequence of alpha-synuclein (Fig. 7). Vaccination resulted in the production of antisera that were capable of recognizing the structure of pathological alpha-synuclein fibrils from multiple synucleinopathies, while not recognizing any linear alpha-synuclein peptides [p < 0.0162] (Fig. 7). Furthermore, each vaccine candidate exhibited a unique pattern of reactivity towards the fibrils contained in the distinct pathologies. The α-SC3-induced antisera showed a significant difference [p < 0.0002] between the synucleinopathy patients and the non-neurologic controls (Fig. 7a). Antisera induced by α-SC6 demonstrated significant differentiation [p < 0.0002] between brain homogenates of dementia with Lewy bodies and Parkinson’s disease patients, however its reactivity against brain homogenates of patients with multiple system atrophy was unable to reach statistical significance with respect to the non-neurologic control samples (Fig. 7b). α-SC8 antisera could significantly differentiate [<0.0001] pathological fibrils derived from brains of patients with dementia with Lewy bodies, but its reactivity for multiple system atrophy and Parkinson’s disease samples did not reach statistical significance in comparison to the non-neurological controls (Fig. 7c). In contrast, the α-SC9 antisera were able to recognize [p < 0.01] pathological fibrils in brain homogenates of dementia with Lewy bodies patients, but did not significantly differentiate between the other pathological groups and non-neurologic controls (Fig. 7d).

**Figure 7:**
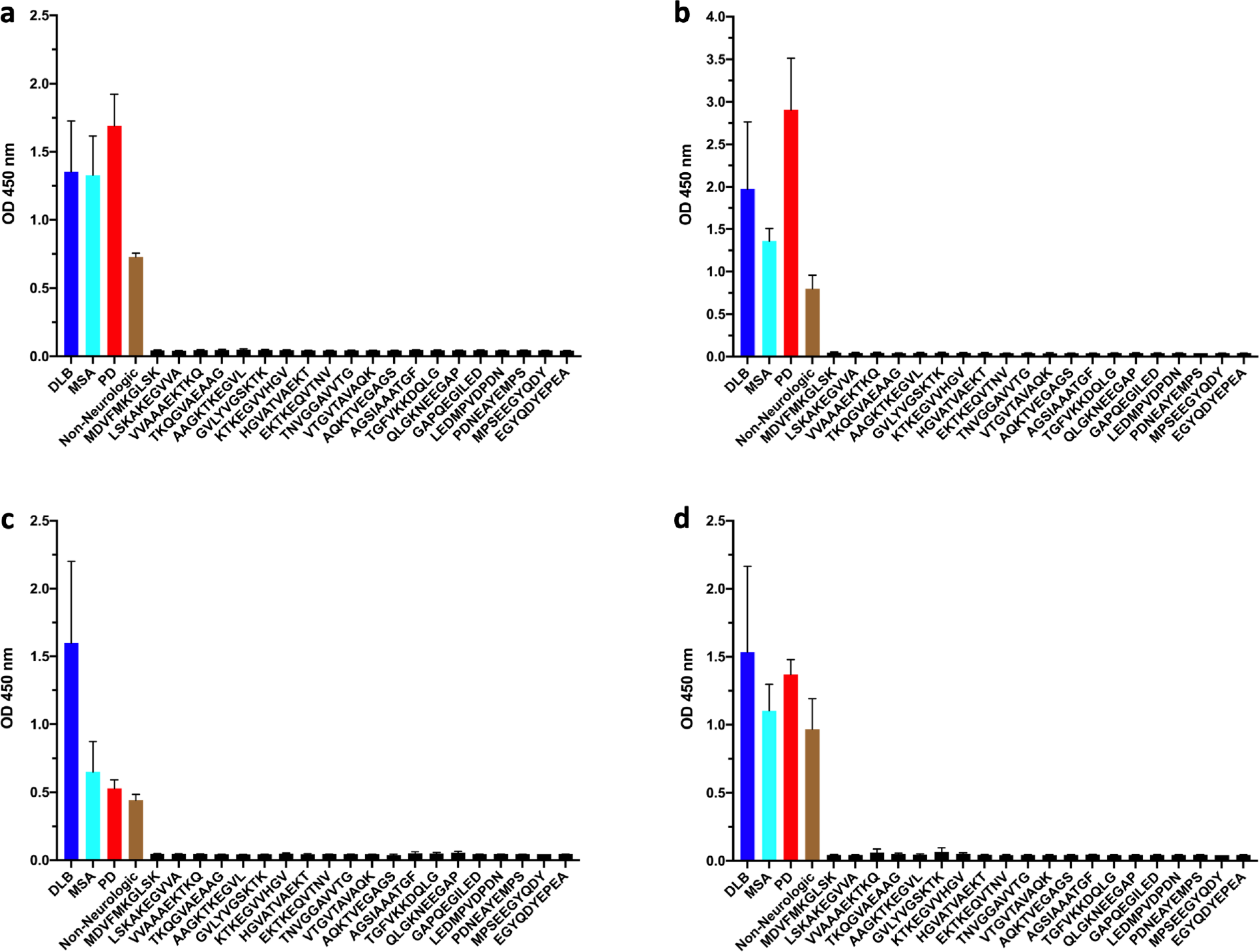
Vaccine candidates mimicking alpha-synuclein fibrils elicit an immune response against conformational epitopes of pathological alpha-synuclein fibrils, instead of linear epitopes. Post-immune antisera derived from the vaccine candidates were tested for their ability to recognize linear peptides corresponding to the entire sequence of alpha-synuclein. As expected, based on the rational structure design, none of the vaccine candidates – α-SC3 (a), α-SC6 (b), α-SC8 (c), α-SC9 (d) – induced an immune response recognizing any linear peptide of alpha-synuclein. In contrast, antisera from the vaccine candidates were able to recognize conformational epitopes present in alpha-synuclein fibrils in human brain lysates obtained from patients with dementia with Lewy bodies (DLB), multiple system atrophy (MSA), and Parkinson’s disease (PD). Data represent the mean value of three human samples per condition with three technical replicates ± SEM. Statistical analysis was performed using one-way ANOVA, followed by Tukey’s post-hoc-test.

**Table 1.** Demographic and clinical profiles of dementia patients and non-neurological controls

| Group | Sex | Age | Diagnosis | Brain region | ID |
| --- | --- | --- | --- | --- | --- |
| Non-neurological | Female | 68 | non-neurological control | cortex | NNC-1 |
|  | Male | 72 | non-neurological control | medulla oblongata | NNC-2 |
|  | Male | 80 | non-neurological control | substantia nigra | NNC-3 |
| DLB | Female | 86 | Dementia with Lewy bodies | caudate with putamen | DLB-1 |
|  | Male | 78 | Dementia with Lewy bodies | caudate with putamen accumbens | DLB-2 |
|  | Male | 72 | Dementia with Lewy bodies | caudate with putamen accumbens | DLB-3 |
| MSA | Female | 61 | Multiple system atrophy | cerebellum | MSA-1 |
|  | Male | 57 | Multiple system atrophy | cerebellum | MSA-2 |
|  | Male | 73 | Multiple system atrophy | cerebellum | MSA-3 |
| PD | Male | 72 | Parkinson's disease | caudate with putamen accumbens | PD-1 |
|  | Male | 82 | Parkinson's disease | caudate with putamen accumbens | PD-2 |
|  | Male | 76 | Parkinson's disease | caudate with putamen accumbens | PD-3 |

In summary, the data demonstrate that post-immune antisera of mice immunized with the vaccine candidates specifically recognize the structural epitopes of alpha-synuclein fibrils, rather than linear epitopes, in a pathological context. Moreover, each candidate showed a unique pattern of reactivity, depending on the disease-associated pathological fibril conformation, which is consistent with the prion-like properties of alpha-synuclein amyloid [57]. Although alpha-synuclein amyloid is not transmissible like prions, evidence suggests the existence of diverse conformers or strains with distinct structural and biochemical properties, different seeding activities, and varying propagation patterns [58], which could contribute to the heterogeneity of neurodegenerative symptoms and disease progression rates observed in synucleinopathies [59].

### Post-immune antisera specifically bind pathological alpha-synuclein in human brain lysates

The ability of the post-immune antisera to recognize to alpha-synuclein in its pathological form was tested by competitive ELISA. Human brain lysates obtained from individuals with different synucleinopathies and non-neurologic controls were used as antigens (Fig. 8). The antibodies in the antisera were first bound to the alpha-synuclein amyloid contained in the brain lysates, limiting their subsequent binding to the vaccine candidates that were used as antigens. The immune specificity of the antibodies to the alpha-synuclein amyloid fibrils present in dementia with Lewy bodies, multiple system atrophy, or Parkinson’s disease samples was measured by a decrease in signal compared to the non-neurologic samples, which were used as a control (Fig. 8).

**Figure 8:**
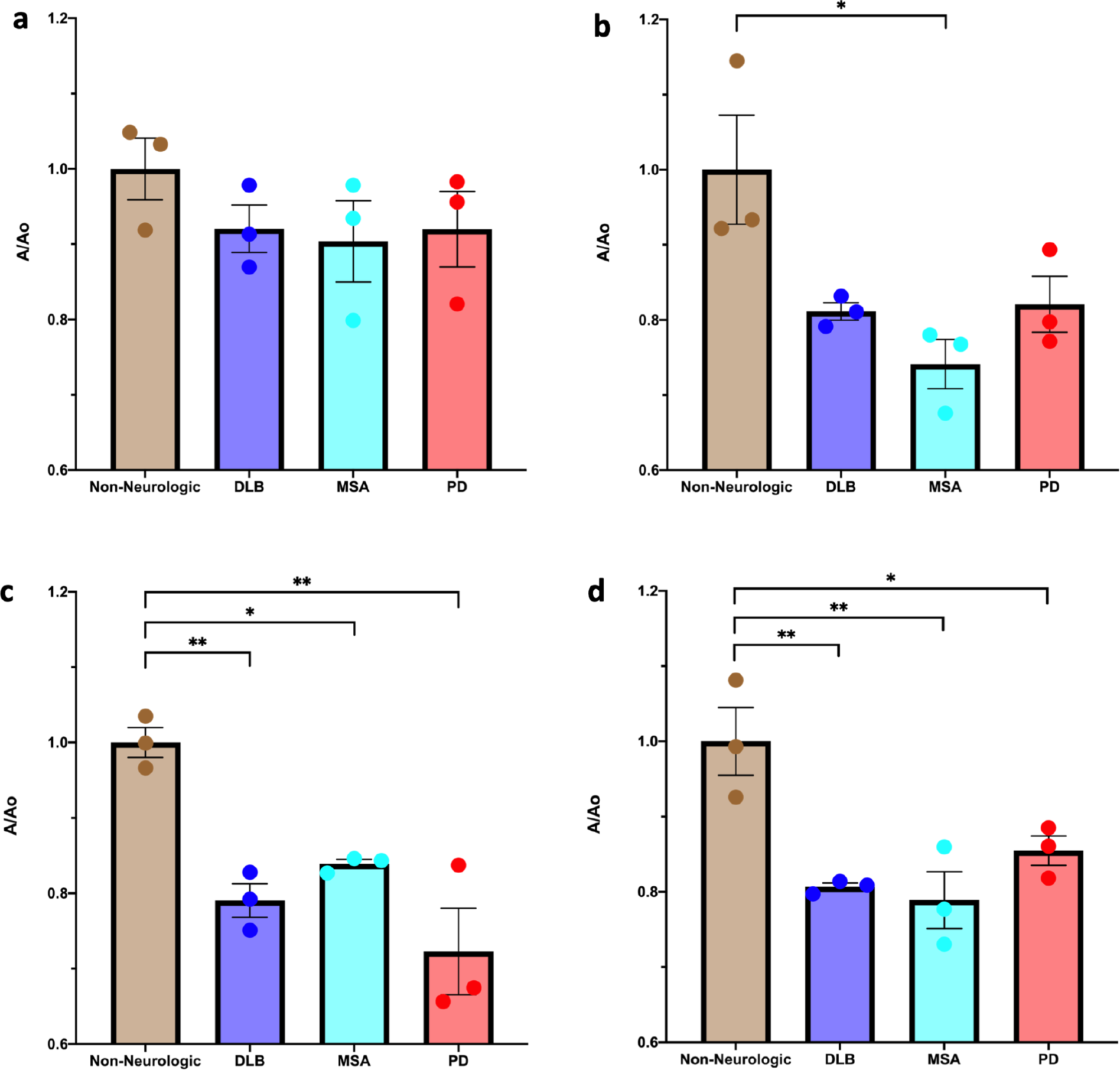
Vaccine candidates elicit immune-specific binding to pathological alpha-synuclein fibrils from patients with synucleinopathies. Post-immune antisera from the synucleinopathy vaccine candidates were tested for their ability to bind to pathological alpha-synuclein fibrils in brain lysates from patients with different synucleinopathies and non-neurologic controls. The antisera-brain-homogenate mixtures were then incubated on plates coated with the original antigen to measure the specificity of the antibodies for the different synucleinopathies. Immune specificity was determined by measuring a decrease in signal relative to the non-neurologic control group. a) α-SC3 antisera did not show any statistical significance compared to the non-neurologic group. b) α-SC6 antisera significantly generated affinity for the multiple system atrophy group compared to the non-neurologic group. c) α-SC8 antisera and d) α-SC9 antisera showed significant affinity for all pathological groups compared to the non-neurologic group, indicating that they specifically recognize the structural epitopes of alpha-synuclein fibrils present in different human synucleinopathies. Data obtained from three independent experiments. The results are expressed as mean values ± SEM for three brain human samples. Statistical analysis was performed using a one-way ANOVA followed by Tukey’s post-test to determine significance [*<0.05, **<0.01].

α-SC3 antisera did not show any significant difference in binding between human brain lysates from disease cases and the non-neurologic controls (Fig. 8a). In the case of the α-SC6 antisera, they showed significantly more binding for pathological fibrils in multiple system atrophy samples [<0.05], however only a trend can be observed for its affinity for dementia with Lewy bodies and Parkinson’s disease compared to the non-neurologic controls (Fig. 8b). Antisera from α-SC8 and α-SC9 immunized mice showed significantly increased binding for all the pathological groups [p < 0.05; < 0.01] in comparison to the non-neurologic group (Fig. 8c,d). Taken together, these results indicate that antisera induced by the vaccine candidates α-SC6, α-SC8, and α-SC9 exhibit specificity for the structural epitopes that come from different human synucleinopathies.

These findings are important because they demonstrate that the candidate vaccines can elicit an immune response that specifically recognizes the pathological forms of alpha-synuclein present in human brain homogenates from patients with synucleinopathies. This suggests that these alpha-synuclein specific vaccines have the potential to target the underlying pathology of these neurodegenerative diseases and could be promising candidates for further development as prophylactic or therapeutic interventions.

### A promising new approach to develop Parkinson’s disease vaccines based on structural insights

There have been numerous attempts to develop active immunotherapies to prevent Parkinson’s disease. The initial attempts involved vaccinating animal models with recombinant human alpha-synuclein protein, resulting in an autoimmune response that affected their physiological functions. Subsequently, among the more promising developments, the two candidate vaccines PD01A and PD03A, based on short peptide sequences that attempt to mimic parts of alpha-synuclein in its native form [38], but not in its aggregated, insoluble form, or in its amyloid fibril form, have failed in clinical trials, showing differences in the specificity of the induced antibodies [40], with a mortality rate of 10% and only 50% of patients developing antibodies against alpha-synuclein [35,44,60]. It is worth noting that similar weaknesses hampered previous attempts to develop vaccines against Alzheimer’s disease, which were based on linear peptides and epitopes without any control of the antigen structure used for immunization, resulting in severe autoimmune reactions in clinical trials [61].

These failures were due to a lack of control over the antigen structure used for immunization and the failure to mimic aggregated alpha-synuclein. Additionally, the structure of alpha-synuclein in its amyloid fibril form was previously unknown. However, atomic-resolution structures of full-length and truncated alpha-synuclein (1-121) fibrils [49,50,62], as well as those obtained from brains published recently [47,63] support the new approach presented in this study for achieving the desired protective immunity against synucleinopathies through the rational design of candidate vaccines based on structure. This novel approach takes into account the secondary and tertiary structure of aggregated alpha-synuclein, as well as the amino acids that are exposed on the surface of alpha-synuclein fibrils, defining specific surface epitopes of alpha-synuclein that are specific for disease. This last step is crucial as it will give our vaccine candidates a unique immunological profile, as shown in this study.

Our study presents a promising new approach for developing vaccines against amyloid diseases, such as Parkinson’s disease and other synucleinopathies, by emphasizing the importance of the disease-relevant pathological conformation of the target, here alpha-synuclein fibrils. Efficacy trials of the candidate vaccines are ongoing in TgM83 mice expressing human alpha-synuclein with the familial A53T mutation. Effective vaccines against alpha-synuclein fibrils are crucial for the prevention and treatment of synucleinopathies, and our approach provides a promising avenue towards achieving this goal.

## Methods

### Selection of vaccine candidate scaffold protein

The innocuous, left-handed beta-solenoid domain of HET-s(218-289) from the filamentous fungus *Podospora anserina* was selected as the scaffold protein from the Protein Data Bank (PDB access code: 2RNM) to introduce exposed amino acids from the surface of alpha-synuclein fibril structures solved by ssNMR (PDB access code: 2N0A) and cryo-EM (PDB access code: 6H6B). The basis for selecting the scaffold protein was a characteristic distance of 4.8 ± 0.2 Å between individual beta-rungs, which is similar to the stacking of alpha-synuclein molecules in their amyloid state [49,50].

### Design and engineering of alpha-synuclein-targeting vaccine candidates

The alpha-synuclein-targeting vaccine candidates were designed using UCSF ChimeraX software. Two different alpha-synuclein structures were used to encompass its fibrillar heterogeneity, one solved by Tuttle et al. using solid-state NMR and the other by Guerrero-Ferreira et al. using cryo-electron microscopy [49,50]. Each vaccine candidate was designed by carefully selecting specific amino acids from the surface of alpha-synuclein fibril structures to be substituted onto the surface of the scaffold protein in a structurally controlled, discontinuous manner to express specific antigenic determinants similar to the alpha-synuclein fibril structures. The repeating nature of the HET-s(218-289) beta-solenoid assembly effectively mimicked the stacking of the parallel in-register beta-sheet structure of the alpha-synuclein fibrils. Out of ten originally designed vaccine candidates, four vaccine candidates (α-SC3, α-SC6, α-SC8, and α-SC9) were successfully expressed and tested.

### Substitution of amino acids for each vaccine candidate

The substitutions of residues for each vaccine candidate targeting synucleinopathies were carefully selected to mimic select epitopes from the structure of two alpha-synuclein molecules per the two rungs of the beta-solenoid scaffold, HET-s(218-289). For α-SC3, the specific amino acids K229Q, D230V, R232N, E234G, and E235G were substituted for the first rung of the scaffold protein to mimic a molecule of alpha-synuclein in its amyloid form, while E265Q, T266V, V268N, and K269G were replaced in the second rung of the scaffold protein to mimic a second molecule of alpha-synuclein that would sit on top of the first one in its amyloid form. Those residues were selected from the ssNMR structure [50]. Meanwhile, for α-SC6, the substitutions were based on the cryo-EM structure reported for alpha-synuclein amyloid fibril, where R232K, E234V, and R236G were substituted for the first rung and V268K, K270V, G271E, and E272G for the second rung [49]. Similarly, for α-SC8, R225K, S227Q, and K229T were substituted for the first rung and T261K, S263Q, and E265T for the second rung, while for α-SC9, S227K, K229V, D230E, and R232A were substituted for the first rung and S263K, E265V, T266E, and V268A for the second rung.

### Cloning and sequencing confirmation of vaccine candidate constructs

The DNA-coding sequences for each engineered vaccine targeting misfolded alpha-synuclein conformers were optimized for expression in *Escherichia coli*. To facilitate cloning, NdeI and BamHI restriction sites were added at the 5’ and 3’ termini of the synthetic genes, which were purchased from BioBasic Inc. (Markham, Ontario). The optimized coding sequences were then digested with the appropriate restriction enzymes at 37 °C for 15 minutes and loaded onto a 1% agarose gel. The digested products were purified and cloned into a pET-21a(+) expression vector.

The pET-21a(+) expression vectors containing the optimized coding sequences were transformed into *E. coli* BL21 (DE3) chemically competent cells by heat shock at 42 °C for 30 s. The transformed cells were recovered by adding Super Optimal broth with Catabolite repression (SOC) media and grown at 37 °C in agitation at 250 rpm for 1 hour. Then, 50 µL of the culture was spread onto pre-warmed plates containing 100 µg/mL ampicillin (Sigma-Aldrich, St. Louis, MO) for antibiotic selection and incubated at 37 °C overnight. Five colonies per construct were isolated and cultured at 37 °C, 250 rpm, overnight in 2YT medium (1.6% tryptone, 1.0% yeast extract and 0.5% NaCl, pH 7.0) containing 100 µg/ml of antibiotic.

DNA plasmids were isolated from the overnight cultures using the QIAprep Spin Miniprep Kit (Qiagen, Hilden, Germany) according to the manufacturer’s instructions. The isolated plasmids were submitted to the Molecular Biology Service Unit (MBSU) of the University of Alberta (Edmonton, AB, Canada) for sequence analysis to confirm the correct insertion of the synthetic genes. Finally, positive colonies were stored in 20% glycerol stock at −80 °C for further protein expression.

### Expression of Vaccine Candidates

A small scrape of bacterial glycerol stock containing the correct DNA coding sequences for each engineered vaccine was used to pre-inoculate 25 mL of 2YT media at 37 °C, 250 rpm, overnight. Then, a 2 L flask containing 500 mL of 2YT media supplemented with ampicillin (100 μg/mL) was inoculated and incubated at 37 °C until an OD600 of ∼0.8 was reached. Following this, the culture was cooled down to 25 °C, and isopropyl beta-D-thiogalactoside (IPTG) was added to a final concentration of 1 mM to induce protein expression overnight at 250 rpm. After induction, the cells were harvested at 5,000 rpm, 4 °C for 25 min and frozen for at least 30 min, and the pellet was resuspended in 20 mL of ice-cold IB buffer (100 mM Tris-HCl, pH 8.0), containing 0.5% Triton X-100, 1 mg/mL lysozyme, and 1X cOmplete™ EDTA-Free Protease Inhibitor Cocktail (Roche, Indianapolis, IN). The resuspended cells were incubated at room temperature (RT) for 30 min and sonicated (Sonifier 250; Branson Ultrasonics, Danbury, CT) for 5 cycles (1 min on and 1 min off) at output voltage 50 with a 50% duty cycle at 4 °C. Subsequently, 3 U/mL benzonase (Merck KGaA, Darmstadt, Germany) per each mL of the original culture were added and the homogenate was incubated for 20 min. The homogenate was centrifuged at 11,000 rpm, 4 °C for 30 min, and the supernatant was then discarded. The resulting pellets, containing the expressed vaccine candidates as inclusion bodies, were frozen for at least 30 min. These pellets were then resuspended in IB buffer, containing 0.5% Triton X-100, 1 mg/mL lysozyme, and 1X cOmplete™ EDTA-Free Protease Inhibitor Cocktail, incubated for 20 min at RT, and sonicated and centrifuged as described previously. This process was repeated between two to five times until the supernatant was clear, then it was finally washed in IB buffer, and the final pellet was centrifuged at 11,000 rpm, 4°C for 30 min. The pellet containing inclusion bodies was then solubilized in 6 M guanidine hydrochloride, 20 mM sodium phosphate, 0.5 M NaCl, pH 8.0, and stirred at RT for 45 min. Subsequently, the homogenate was clarified by ultracentrifugation at 45,000 rpm for 35 min at 4°C for further purification.

### Vaccine-candidate purification

The recombinant protein of each vaccine candidate was purified under denaturing conditions using a biologic duoflow™ chromatography system (Bio-Rad Laboratories, Hercules, USA). A HisTrap™ HP column (GE Healthcare Life Sciences, Piscataway, NJ, USA) was equilibrated with equilibrium buffer (8 M Urea, 20 mM Sodium Phosphate, 0.5 M NaCl, 10 mM imidazole, pH 8.0) to prepare the column for sample loading. Once the absorbance at 280 nm reached a stable baseline measurement, the clarified lysate was loaded onto the column, and the column was washed with the same equilibrium buffer until the absorbance at 280 nm again reached a stable baseline measurement to remove unbound proteins. A linear gradient from 10 mM to 500 mM imidazole (the other components were the same as the equilibrium buffer) was used to elute the bound protein, and the fractions containing the recombinant vaccine candidates were automatically collected using BioFrac™ Fraction Collector (Bio-Rad Laboratories, Hercules, USA) when the 280 nm absorbance was greater than 0.05. The purity and identity of the eluted proteins were confirmed by SDS-PAGE.

### Vaccine candidates fibrillization process

Following purification, the vaccine candidates were subjected to a buffer exchange process to remove any residual salts or other impurities that could interfere with subsequent experiments. The HiTrap® Desalting Columns (GE Healthcare Life Sciences, Piscataway, NJ, USA) were used to exchange the buffer, and a low pH of approximately 2.8 (175 mM acetic acid) was chosen to help maintain the stability of the protein. The desalted protein samples were then subjected to an in-vitro fibrillization process to encourage the formation of amyloid fibrils, which are the primary vaccine candidates. The pH was increased to 7.5 using 3 M Tris base (∼pH 13.0), and the samples were stirred at 600 rpm for a minimum of 5 days at RT.

### Protein electrophoresis

The purity and molecular weight of the vaccine candidates were evaluated using sodium dodecyl sulfate–polyacrylamide gel electrophoresis (SDS-PAGE). To prepare the samples, 5x SDS sample buffer containing beta-mercaptoethanol was added to the purified vaccine candidates, which were then heated at 95 °C for 5 min. The samples were then loaded onto NuPAGE 12% Bis-Tris protein gels (Thermo Fisher Scientific, Waltham, MA) and electrophoresed at 170 V for 40 min. Gels were stained with Bio-safe Coomassie stain (Bio-Rad Laboratories, Hercules, CA) and visualized. The molecular weight of the vaccine candidates was estimated by comparing the migration of the protein bands to the molecular weight markers. The accuracy of the molecular weight determination was confirmed using ExPASY’s ProtParam tool.

### Evaluation of Structural Stability

To evaluate the secondary structures of the vaccine candidates, Far-UV circular dichroism spectra were recorded using a Chirascan™ Circular Dichroism (CD) Spectrometer system. Amyloid fibrils were buffer exchanged in 10 mM sodium phosphate/tris buffer at pH 7.5 and 0.2–0.5 mg/mL of sample protein was used for the assay. CD spectra were recorded at 25 °C in a 0.1 cm cell from 190 to 260 nm by 1.0 nm increments. A buffer CD spectrum was subtracted from the protein CD spectra. Each spectrum was generated from the average of 10 scans.

### Transmission electron microscopy

Negative stain transmission electron microscopy was used to confirm the ability of the vaccine candidates to self-assemble into the typical amyloid structure of the native HET-s(218-289) protein and to confirm fibril formation. Carbon-coated copper grids with a mesh size of 200 squares were glow discharged at 15 mA, 0.39 mbar for 1 min. An aliquot of 5 µL of purified sample was loaded and absorbed onto the grid for 1 minute and washed with two drops (50 µL) of filtered ammonium acetate: one drop of 100 mM and the second of 10 mM. The grid was then stained by adding 10 µL of 2% filtered uranyl acetate (Electron Microscopy Sciences, Hatfield, PA). Grids were then blotted dry with filter paper and stored at RT and visualized by using a bottom-mounted Eagle 4k x 4k camera on a Tecnai F20 TEM (FEI Company, Hillsboro, OR) operating at 200 kV.

### Animals

The animal study was conducted in accordance with the National Institute of Health’s “Guide for the Care and Use of Laboratory Animals” and the Canadian Council on Animal Care. All procedures were approved by the Animal Care and Use Committee at the University of Alberta. Mice were housed in a temperature-controlled room (22 °C) with a 12 h light/dark cycle and had ad libitum access to food and water. The animals were handled with care throughout the study to minimize discomfort and distress. FVB mice were used as the wild type. The hemizygous presence of the human A53T alpha-synuclein transgene in TgM83^+/−^ (B6;C3-Tg(*Prnp*-SNCA*A53T)83Vle/J) mice was confirmed by real-time PCR analysis.

### Optimization of Immunization Regimen

To determine the optimal immunization dose, different priming doses of the vaccine candidate (100, 50, 25, or 5 μg) were administered intraperitoneally to both male and female mice aged eight weeks with a body weight of approximately 22 g. The vaccine candidate was sonicated in 50 µL of PBS and emulsified with 50 µL of Freund’s complete adjuvant (Sigma-Aldrich, St. Louis, MO). This was followed by three biweekly boosters of half the priming dose, combined with Freund’s incomplete adjuvant in a 50/50 (v/v) ratio. Enzyme-linked immunosorbent assay (ELISA) was used to determine the post-immune antisera titers. Blood was collected before the start of immunization (pre-immune sera) and every two weeks before each boost, with post-immune antisera collected two weeks after the final boost. Once the optimal immunization regime was determined, all animal immunizations were performed at that prime and boost dose.

### Indirect ELISA

High-binding microplate strips (Santa Cruz Biotechnology Inc., Santa Cruz, CA) were used for the ELISA. The strips were coated with 0.5 µg of each sonicated vaccine candidate, diluted in coating buffer (PBS), and incubated overnight at 4 °C. The strips were then washed with TBST (0.1% Tween 20) and blocked with 3% BSA in TBST (0.05% Tween 20) for 2 hours at RT. After blocking, the primary antibody (post-immune sera) was added to each well, serially diluted in blocking buffer, and incubated overnight at 4 °C. The plate was washed again with TBST, and a secondary HRP-goat anti-mouse, diluted 1:5000 in 3% BSA TBS, was added to each well and incubated for 1 hour at RT. Following another wash step, 3,3’,5,5’-Tetramethylbenzidine (TMB) substrate solution (SurModics, Eden Prairie, MN) was added to each well and incubated for 5–30 minutes at RT until the color developed. Finally, the reaction was stopped by adding stop solution (2 N H_2_SO_4_) to each well, and the absorbance of each well was measured at 450 nm using a microplate reader. Each wash step consisted of 3 washes. For experiments involving human brain homogenates or alpha-synuclein peptides to coat the strips, 4 µg of lysates or 40 µg of alpha-synuclein peptides were added to each well.

### Competitive ELISA

Post-immune antisera were added to sonicated human brain lysates containing alpha-synuclein amyloid fibrils, which were used as antigens in non-binding microplates (Greiner Bio-One, Kremsmünster, Austria). The mixture was incubated for 1 hour with agitation at RT and then transferred to high-binding microplate strips (Santa Cruz Biotechnology Inc., Santa Cruz, CA) previously coated with 0.5 µg of each sonicated vaccine candidate in coating buffer (PBS), following the indirect ELISA procedure. The strips were washed, blocked, and developed as in the indirect ELISA. Antibody binding to alpha-synuclein amyloid fibrils present in the different patient brain homogenate samples was measured by a decrease in signal from the non-neurologic group, which was measured at 450 nm.

### Preparation of human and mouse brain lysates

Human 10% (w/v) brain homogenates in PBS were prepared from brain tissues of patients who died with dementia with Lewy bodies, multiple system atrophy, or Parkinson’s disease, and of control individuals who died due to non-neurologic causes. The protein concentration of each brain homogenate was quantified using the Bradford assay. Brain homogenates of tissues from hemizygous TgM83^+/−^ mice that were intracerebrally challenged with preformed alpha-synuclein fibrils or with BSA were prepared in the same way.

### Statistical analysis

All data were expressed as mean ± standard error of triplicate measurements. The normality of the data was assessed using the Shapiro-Wilk test, and as the data was normally distributed (p < 0.05). A one-way ANOVA test followed by Tukey’s post-hoc test was used to identify significant differences between groups using GraphPad Prism 8.0.1 software (GraphPad Software, San Diego, CA, USA). A p-value below 0.05 was considered statistically significant.

## Acknowledgements

We gratefully acknowledge support from the Weston Brain Institute (RR191051 to H.W. and G.T.) and the Michael J. Fox Foundation for Parkinson’s Research (MJFF-009841 to G.T. and H.W.). We furthermore acknowledge the receipt of valuable human brain tissue samples from Dr. Inga Zerr at the Clinical Dementia Center, Georg-August University in Göttingen, Germany. Additional brain samples were obtained from The Netherlands Brain Bank, Netherlands Institute for Neuroscience, Amsterdam (open access: www.brainbank.nl). All Material has been collected from donors for or from whom a written informed consent for a brain autopsy and the use of the material and clinical information for research purposes had been.

---

**Figure S1:**
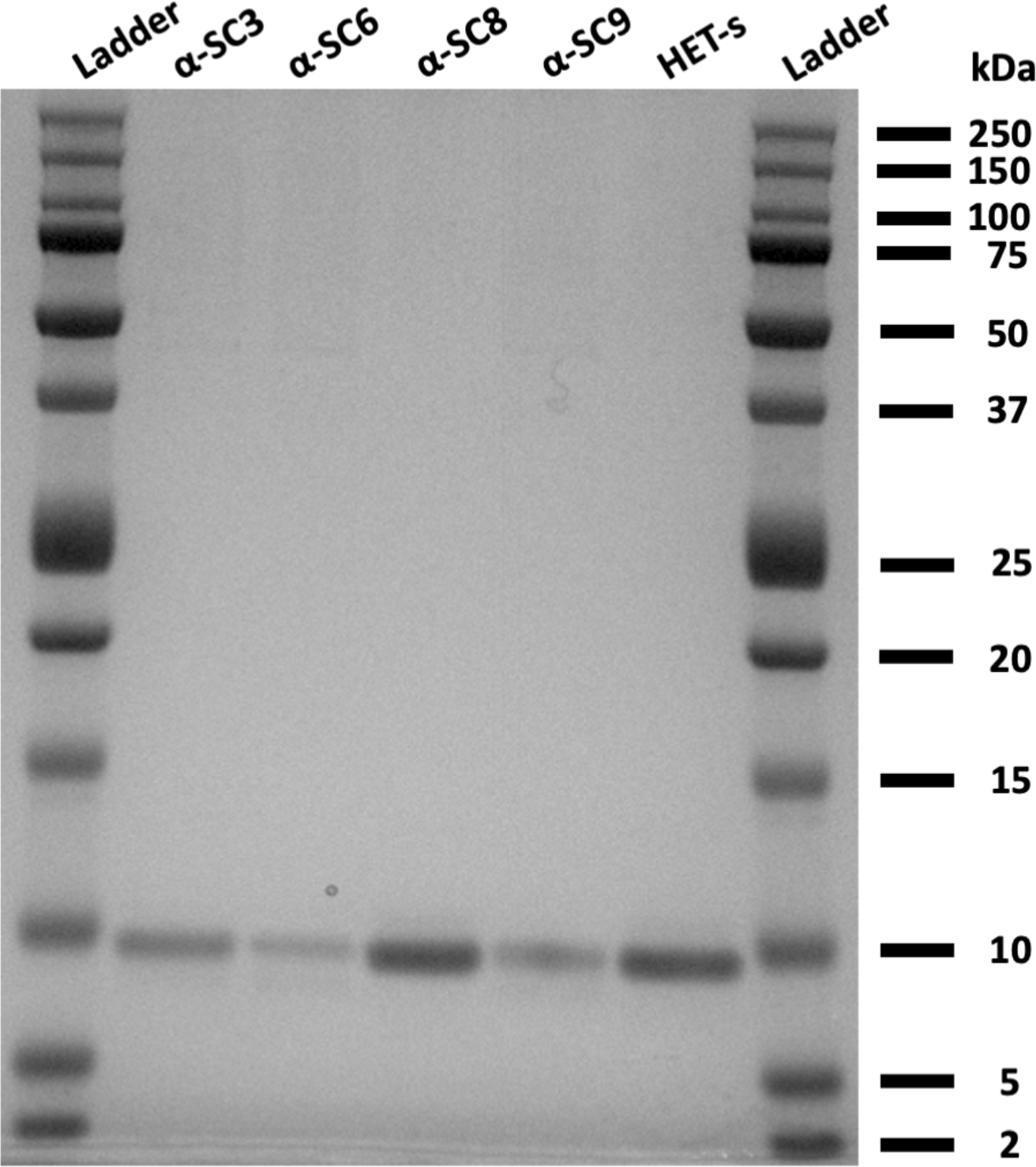
Successful production and purification of alpha-synuclein-targeting vaccine candidates for further study. The engineered proteins were optimized for expression in *Escherichia coli*, purified by affinity chromatography, and desalted. The resulting proteins were resolved on a NuPAGE 12% Bis-Tris protein gel, and found to have a molecular mass not significantly different from the original scaffold protein.

## Notes

### Competing Interest Statement

The authors have declared no competing interest.

